# Cell wall integrity signaling regulates cell wall regeneration via transcriptional activation in *Chlamydomonas reinhardtii*

**DOI:** 10.1101/543280

**Authors:** Evan Cronmiller, Deepak Toor, Nai Chun Shao, Thamali Kariyawasam, Ming Hsiu Wang, Jae-Hyeok Lee

**Affiliations:** Department of Botany, University of British Columbia 6270 University Blvd., Vancouver, Canada

**Keywords:** cell wall, cell wall integrity, gametolysin, Chlamydomonas, transcriptional activation, osmotic stress, cell wall defective mutant

## Abstract

An intact cell wall is critical for protecting the cell from osmotic challenges and harmful environments. Signaling mechanisms are necessary to monitor cell wall integrity and to regulate cell wall production and remodeling during growth and division cycles. The green alga, *Chlamydomonas*, has a proteinaceous cell wall of defined structure that is readily removed by gametolysin (g-lysin), a metalloprotease released during sexual mating. Naked cells treated with g-lysin induce the mRNA accumulation of > 100 cell wall-related genes within an hour, offering a system to study signaling and regulatory mechanisms for *de novo* cell wall assembly. Combining quantitative RT-PCR and luciferase reporter assays to probe transcript accumulation and promoter activity, we revealed that up to 500-fold upregulation of cell wall-related genes was driven at least partly by transcriptional activation upon g-lysin treatment. To investigate how naked cells trigger this rapid transcriptional activation, we tested whether osmotic stress and cell wall integrity are involved in this process. Under a constant hypotonic condition, comparable levels of cell wall-gene activation were observed by g-lysin treatment. In contrast, cells in an iso- or hypertonic condition showed up to 80% reduction in the g-lysin-induced gene activation, suggesting that hypotonic conditions are required for full-scale responses to g-lysin treatment. To test whether mechanical perturbation is involved, we isolated and examined a new set of cell wall mutants with defective or little cell walls. All cell wall mutants examined showed a constitutive upregulation of cell wall-related genes at the level, which would only be achieved by the g-lysin treatment in wild-type cells. Our study suggests a signaling that monitors mechanical defects of cell walls and regulates cell wall-gene expression in *Chlamydomonas*, which may relate to cell wall integrity signaling mechanisms in plants.

## Introduction

The cell wall is an extracellular compartment that surrounds cells and provides protection and buffering from environmental fluctuation. The single-celled alga *Chlamydomonas reinhardtii* constantly builds and modifies its cell walls throughout its life cycle^1^. Occasionally, when two nitrogen-starved sexual gametes encounter each other, they initiate a ‘mating reaction’ and remove their cell walls in preparation for cell fusion and subsequent zygotic wall assembly^2^. Consequently, the cells become naked and exposed to their environment and immediately rebuild their cell walls. A failure to do so may lyse the cells under the hypotonic freshwater environments where *C. reinhardtii* live. Given this importance of cell wall regeneration, in this study we investigated how cells sense ‘nakedness’ to rebuild their walls.

The morphology and composition of cell walls vary across Viridiplantae comprised of green algae and plants^3,4^. The vegetative cell wall of dividing cells of a green alga, *Chlamydomonas reinhardtii* is made almost entirely of proteins, including hydroxyproline (Hyp)-rich glycoproteins, and its multi-layered architecture makes it both hardy and flexible^5,6,7^. This architecture can accommodate a ten-fold increase in cell size during the light phase of the daily light/dark cycle. *C. reinhardtii* cells build a second type of cell wall during zygote development following the mating between *plus* and *minus* sexual gametes^8,9^. The mating reaction leads to the release of a metalloprotease, gametolysin (g-lysin), which sheds the cell wall to allow gamete fusion and subsequent *de novo* assembly of a strong zygote cell wall^2,10^. This zygote wall is chemical-resistant and desiccation-tolerant, providing a safe environment for the zygotes to lay dormant until conditions are once again favorable^11,12,13^.

Of the cell wall structural components, many Hyp-rich glycoprotein-encoding genes are upregulated as early as 15 minutes after cell wall shedding by g-lysin treatment^14,15,16^. Hoffmann and Beck^17^ examined in detail the regulation of three gamete-specific (GAS) Hyp-rich pherophorin-encoding genes, *GAS28*, *GAS30*, and *GAS31*, whose expression increases upon g-lysin treatment without responding to variable osmotic conditions. This study suggested that cell wall removal upregulates *GAS* gene expression. It remains unknown how cell wall removal upregulates these three gamete-specific gene transcripts or whether their finding for these GAS genes is applicable to the other g-lysin-inducible cell wall-related genes.

The importance of signaling triggered by g-lysin treatment is suggested by the number of genes regulated by this signal. A recent study using transcriptome analysis revealed 143 genes up-regulated within one hour following g-lysin treatment^18^, suggesting that a signal triggered by g-lysin treatment may control the assembly of the vegetative cell wall. Comparative analysis of this g-lysin-induced transcriptome with an early zygote transcriptome identified two subsets of genes in the g-lysin-induced transcriptome, distinguished by the presence or absence of upregulation in early zygotes^19^. The latter, the vegetative wall-specific g-lysin-induced gene subset (C24 or gL-EZ^19^) includes 36 Hyp-rich glycoprotein-encoding genes particularly enriched in the pherophorin family, likely specific for the vegetative wall structure. The other subset, which comprises genes common to both vegetative and zygotic walls (C44 or gL+EZ^19^), includes 67 genes involved in protein glycosylation and protein secretion, indicating that g-lysin-induced cell wall removal indeed controls cell wall assembly together with the upregulation of structural cell wall protein genes. Hereafter, we refer to these two subsets of cell wall-related genes as CW genes of the protein processing type and the structural protein type.

Here, we present mechanistic insights into the elusive signal generated by g-lysin-induced cell wall removal as a critical step forward from the pioneering study by Hoffmann and Beck^17^. First, we examined whether CW genes are activated via transcriptional and post-transcriptional mechanisms using our promoter-reporter transgenic strains. Second, we evaluated three signals, osmotic stress, the release of digested cell wall fragments, and the loss in cell wall integrity – expected during g-lysin treatment – as potential triggers for the activation of CW genes using cell wall defective (*cwd*) mutants that we isolated and reported in this study (see Supplementary Figure S1 for the illustrations of three hypothetical signals).

Our data show that osmotic stress plays a minor role for fully activating the cell wall regeneration, whereas compromised cell wall integrity plays a dominant role in this process regardless of the g-lysin treatment. Taken together, we propose a new signaling mechanism that senses detached walls or unassembled wall fragment as defective cell walls in *C. reinhardtii*. The same signal may also control cell wall assembly in *C. reinhardtii* and other algae for commercial biomass production. Placing our promoter-reporter system in *cwd* mutants that exhibit constitutive CW gene expression will provide an excellent tool for a suppressor screen where molecular components of the cell wall integrity signaling can be identified.

## Results

### g-lysin treatment induces transcriptional activation of CW genes

A majority of the genes in the CW gene sets fell into two functional groups; protein processing (i.e., related to translation or glycosylation) and structural cell wall proteins. To survey the CW genes systematically, we selected three genes related to protein processing; *SEC61G*, *AraGT1*, and *RHM1*, and four structural cell wall proteins; *GAS28*, *GAS30*, *GAS31*, and *PHC19* (Table 1). According to the reads per million kilobases mapped (RPKM) values as proxy to absolute expression level, taken from the Ning et al.^18^ dataset, the delta RPKM values between the g-lysin-untreated and g-lysin-treated cells for the selected genes indicate a substantial increase in gene expression within 1 hour upon g-lysin treatment (Table 1).

**Table 1.**
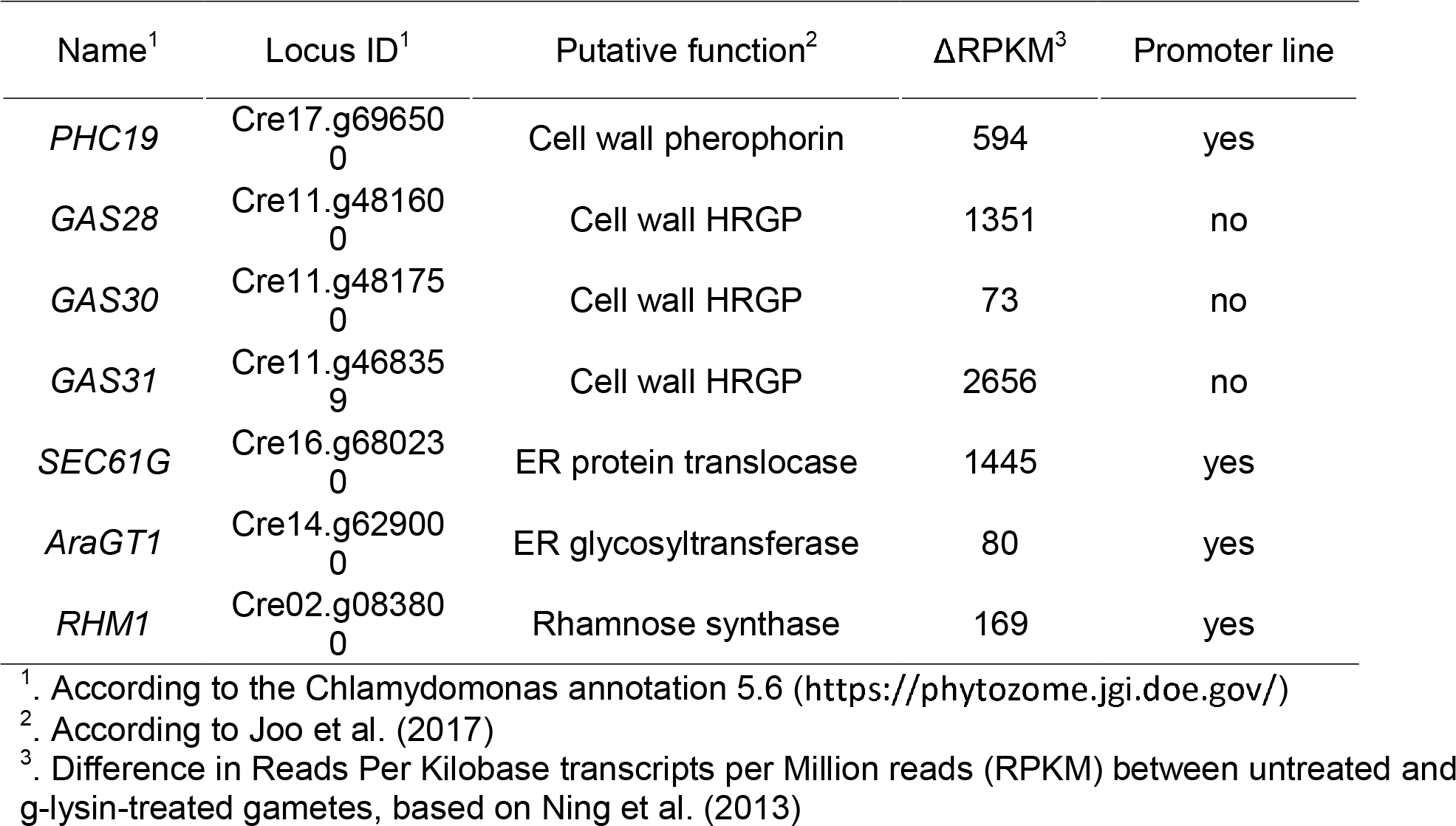
Genes of interest selected from curated transcriptome data.

To confirm the reported transcriptome results and quantify the g-lysin induced gene expression, we analyzed the expression of our selected genes by reverse transcription and quantitative PCR (RT-qPCR) in gamete cells where little growth-related cell wall remodeling is expected. Following the treatment of gametic cells with g-lysin, a significant change in expression was observed for the structural protein genes, ranging between 145- and 508-fold increases (Figure 1a). The protein processing genes showed modest increases between four- and 33-fold (Figure 1b).

**Figure 1.**
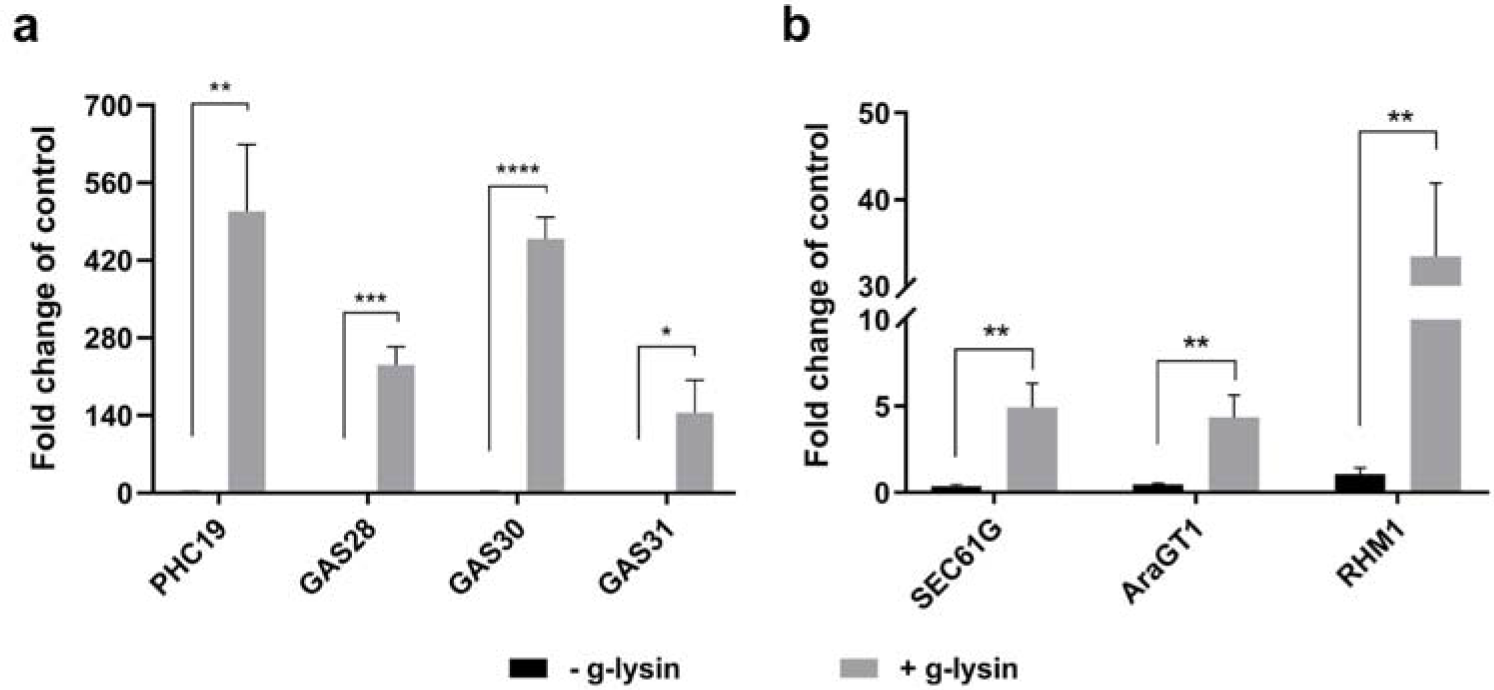
Transcription of CW genes in response to gametolysin treatment shows global upregulation. (**a-b**) Bar graphs represent the change in expression for cell wall protein-encoding (a) and protein processing-related (b) genes in wild type (CC-125) cells. Untreated control samples are represented by black bars, grey bars represent cells treated with g-lysin. Gene expression is quantified in terms of fold change compared to the samples before the g-lysin treatment. Error bars represent one standard deviation from the mean of biological triplicate samples. Welch’s t-test indicates statistical significance at p ≤ 0.05 (*), p ≤ 0.01 (**), p ≤ 0.001 (***), p ≤ 0.0001 (****); ⍰ = 0.05.

To distinguish the transcriptional and the post-transcriptional mechanisms for the upregulation by g-lysin treatment, transgenic strains harboring promoter-luciferase constructs were used to probe promoter activities for *SEC61G*, *AraGT1*, *RHM1*, and *PHC19* genes, as described in our previous study^19^. Two independent transgenic lines for each promoter-reporter construct showed a definite increase, ranging between four- and 210-fold in luciferase expression in response to g-lysin treatment (Figure 2). This result suggests that most of the cell wall-regulated genes are activated at the transcriptional level. Note that the increase of promoter activity for a given gene did not always match the extent to which the transcript accumulated in response to g-lysin. For example, the *SEC61G* promoter showed a hundreds-fold increase in activity when treated with g-lysin, while the mRNA levels increased only modestly (Figure 1). This discrepancy may be due to the post-transcriptional regulation of *SEC61G* expression.

**Figure 2.**
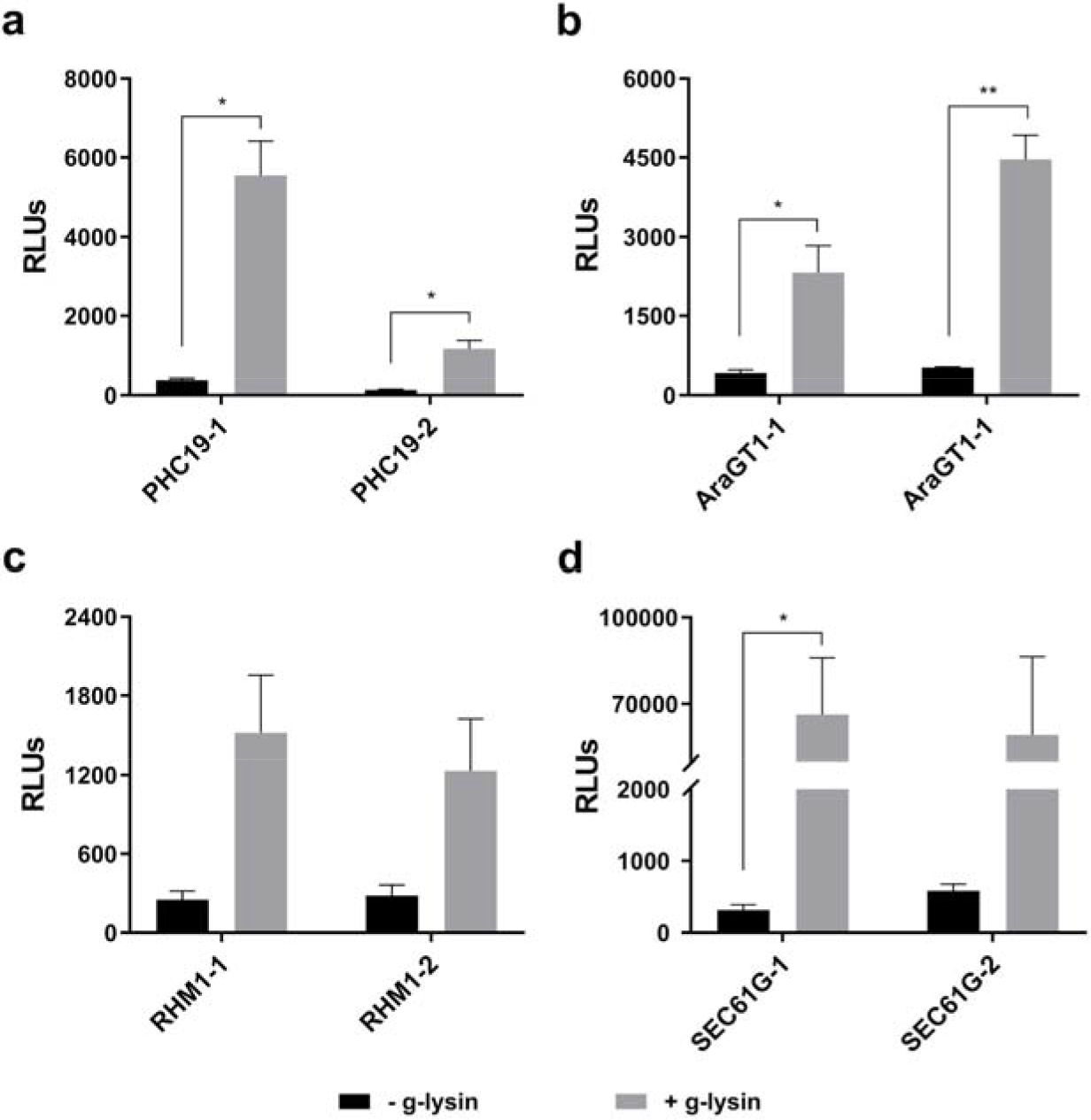
Promoter-driven luciferase activity increases in response to g-lysin treatment. Bar graphs represent the change in promoter activity of *PHC19* (a), *AraGT1* (b), *RHM1* (c), and *SEC61G* (d) genes, using two independent promoter-transformed lines. Luciferase activity is expressed as relative light units (RLUs) based on luminescence quantification. Untreated gamete control samples are represented by black bars. Grey bars represent gamete cells treated with g-lysin. Error bars are one standard deviation from the mean of biological triplicate data. Welch’s t-test indicates statistical significance at p ≤ 0.05 (*), p ≤ 0.01 (**); α = 0.05.

### Translational inhibition revealed the complex regulatory network of the g-lysin-induced CW gene expression

One of the critical features of signaling pathways is a hierarchical structure, where a primary response leads to a secondary response, dependent on the protein produced from the primary response. To examine such a hierarchy, we examined g-lysin-induced transcript accumulation in cells treated with the eukaryotic protein synthesis inhibitor cycloheximide (CHX). The CHX treatment itself did not affect much of the CW gene expression, showing less than a three-fold change (Figure 3). By comparing the g-lysin-induced gene expression between CHX-treated and untreated cells, we identified three patterns. First, *RHM1* and *PHC19* showed two-to five-fold reduction of the g-lysin-induced up-regulation but remained responsive to g-lysin treatment (Figure 3). Second, *GAS28* and *GAS30* showed near complete inhibition (< four-fold) of the g-lysin-induced up-regulation in CHX-treated cells, indicating that *GAS28* and *GAS30* are regulated by a secondary response, which is consistent with a previous report^17^. Third, *AraGT1* and *GAS31* showed three- and eight-fold increases in the g-lysin-induced up-regulation when pretreated with CHX. This increase suggests that some of the early responsive CW genes such as *AraGT1* and *GAS31* are down-regulated as negative feedback by a regulator either short-lived or up-regulated by the g-lysin treatment. Overall, our result showed that *AraGT1*, *RHM1*, *PHC19* and *GAS31* are primary targets and *GAS28* and *GAS30* are secondary targets of the g-lysin-induced signaling pathway.

**Figure 3.**
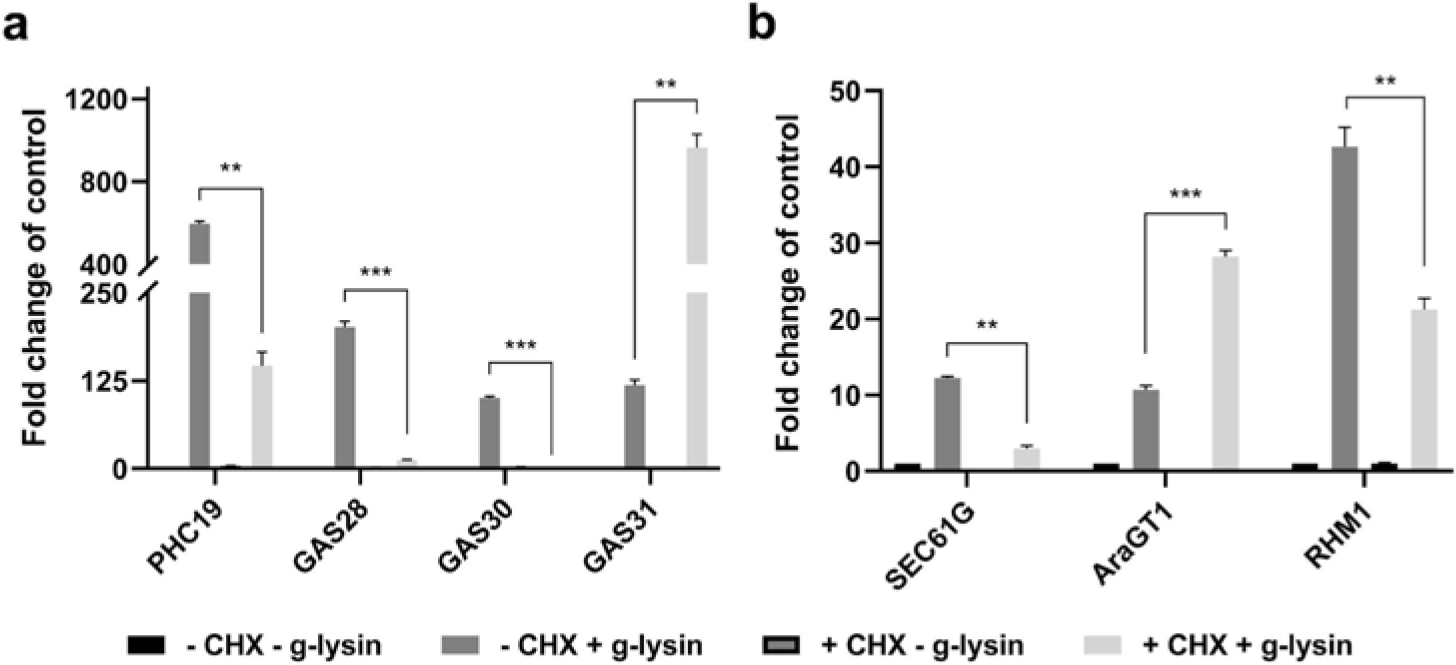
Transcripts are differentially affected by CHX pretreatment prior to gametolysin treatment. (**a-b**) Bar graphs represent the change in gene expression for cell wall protein (a) and protein processing type (b) CW gene transcripts in wild type (CC-125) cells. Untreated gamete control samples represented by black bars, -CHX -g-lysin; medium grey bars for CHX treated cells, +CHX; dark grey bars for cells treated with g-lysin, +g-lysin; light grey bars for both CHX and g-lysin-treated cells, +CHX +g-lysin. Gene expression is quantified in terms of fold change of the untreated control condition. Error bars represent one standard deviation from the mean of biological duplicate samples. Welch’s t-test indicates statistical significance at p ≤ 0.01 (**), p ≤ 0.001 (***); α = 0.05.

### Osmotic stress does not initiate the signal that induces CW gene activation

In *C. reinhardtii*, osmotic balance is controlled by the combined action of a pair of contractile vacuoles (CVs), which pump excess water out of the cell, and the cell wall, which protects the cell from lysing in under the naturally hypotonic freshwater environments where *C. reinhardtii* live (Supplementary Text S1 for details). Therefore, when cells lose their cell walls, osmotic stresses are likely to be incurred on the cells. The effects of changing osmotic conditions can be examined by using the CV cycle as a proxy for water flux.

To assess the effect of osmotic stress on CW gene expression, we analyzed cells transferred from Tris-acetate-phosphate (TAP), a standard growth medium (64 mOsm/L) to half-diluted TAP (1/2 TAP, 32 mOsm/L), where the CV cycle shortens, and sorbitol-supplemented hypertonic condition (TAP+SS, 204 mOsm/L), where the CV cycle stops (Supplemental Table S1). The structural protein genes showed two-to four-fold upregulation when adjusted to the hypertonic condition (TAP+SS), whereas only non-significant changes were observed in the strong hypotonic condition (½ TAP) (Supplementary Figure S2). *SEC61G* showed no significant difference under osmotic stress. Overall, the level of up-regulation observed under osmotic stress is, however, far smaller than the fold-induction following g-lysin treatment, suggesting only a minor role in CW gene regulation.

It may be possible that the cellular osmotic state rather than osmotic stress may be necessary for the cells to activate CW genes. To test this possibility, we designed an experiment that puts cells in different osmotic conditions while reducing osmotic stress to a minimum during g-lysin treatment. Cells were first adapted to hypo-(standard TAP, 64 mOsm/L), iso-(175 mOsm/L), and hypertonic (204 mOsm/L) conditions according to our CV cycle observations. The preconditioned cells were then treated with g-lysin prepared in media of the same osmolarity (see methods for details). In the iso- and hypertonic conditions, we observed a drastic loss, between 72% and 92%, of the fold-induction observed for the CW genes upon g-lysin treatment (Figure 4). This result suggests that a hypotonic condition or contractile vacuole cycling is critical for *C. reinhardtii* cells to fully activate the CW genes via the g-lysin-induced signaling pathway.

**Figure 4.**
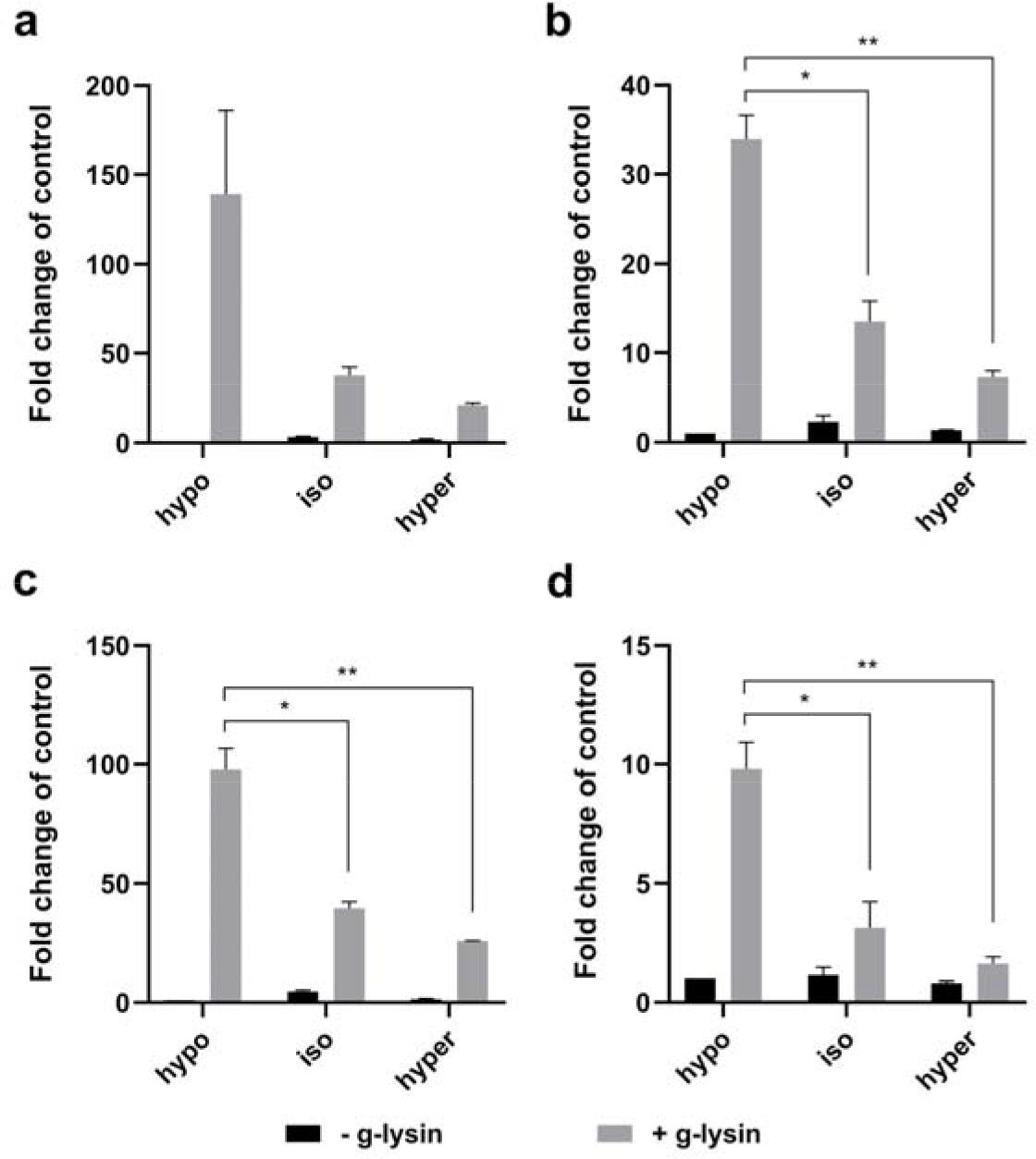
Isotonic and hypertonic conditions negatively affect the g-lysin-induced CW gene activation. (**a-d**) Bar graphs represent the change in gene expression by g-lysin treatment for *PHC19* (a), *GAS28* (b), *GAS30* (c), and *SEC61G* (d) in wild type (CC-125) cells. g-lysin treatment was done in three constant osmotic conditions, hypo-(64 mOsm), iso-(175 mOsm), and hyper-(204 mOsm). Untreated gamete control samples represented by black bars, -g-lysin; medium grey bars for g-lysin-treated cells, +g-lysin. Gene expression is quantified in terms of fold change of the untreated control in hypotonic condition. Error bars represent one standard deviation from the mean of biological duplicate samples. Welch’s t-test indicates statistical significance at p ≤ 0.05 (*), p ≤ 0.01 (**); α = 0.05.

### Mechanical perturbation of cell walls triggers the signal for CW gene activation

Our results discovered the critical importance of natural osmotic conditions for the full-scale activation of CW genes. Nonetheless, CW gene activation was not completely abolished even in the absence of contractile vacuole cycling, thus the trigger of CW gene activation remained to be determined. Thereby, we investigated the mechanical integrity of the cell wall as the potential trigger of CW gene activation. The cell wall integrity was examined by testing whether the cells lyse in the presence of 0.1% non-ionic detergent NP-40 (or Tergitol) – a substance to which cells with fully intact cell walls are undisturbed, while membranes of cells with defective cell walls are compromised.

Earlier studies of cell wall-defective mutants exhibiting lysis upon NP-40 treatment categorized mutant phenotypes into three distinct groups: A) cells producing normal-looking walls attached to the plasma membrane, B) cells producing walls but not connected to the plasma membrane, and C) cells producing minute amounts of wall material^20,21,22^. These groups represent the three sequential consequences caused by g-lysin treatment: cracking the wall integrity, detaching the cell wall from the plasma membrane, and complete removal of the wall. Therefore, whether CW genes are up-regulated without g-lysin treatment in cell wall defective mutants would inform about the involvement of cell wall integrity signaling for CW gene activation caused by the g-lysin treatment.

Most of the cell wall defective strains frequently used in *C. reinhardtii* research were isolated in the early 70s. Therefore, the available cell wall defective strains may have accumulated spontaneous mutations during their long-term culture. To assess the cell wall defective conditions without complex strain history, we isolated a new set of cell wall defective mutants based on their sensitivity to 0.1% NP-40. We selected a subset of these mutants based on NP-40 sensitivity and cell wall morphology to represent diverse mutant types. Our collection included four mutants (*cwd1-4*) from our mutant library and one historical mutant, *cw15*, whose phenotypes are summarized in Table 2. In our collection, *cwd1* and *cw15* contain no residual wall mutant based on their fully round shape and the absence of hollow in the phase-contrast images; *cwd2* and *cwd3* exhibit abnormal cell wall morphologies; and *cwd4* is a putative cell wall detachment mutant (Figure 5). *cw15*, *cwd1*, *cwd3*, and *cwd4* showed high sensitivity to NP-40 (>90% cells burst within 2 min.) while *cwd2* showed medium sensitivity (70-90% cells burst).

**Table 2.** Summary of selected cell wall defective mutants.

| Cell line | Cell morphology | Cell wall phenotype | cw<br>classification<sup>1</sup> | NP-40<br>sensitivity |
| --- | --- | --- | --- | --- |
| *cw15*<br>(CC-3491) | round cells | some material in<br>medium | C | high |
| *cwd1* | round cells | no wall material<br>present | C | high |
| *cwd2* | teardrop shaped;<br>cell aggregates | unevenly distributed | B | medium |
| *cwd3* | oval cells | ruffled-looking<br>edges | B | high |
| *cwd4* | gap between<br>membrane and cell wall | normal looking | B | high |

<sup>1</sup>. (A) cells producing normal-looking walls attached to the plasma membrane, (B) cells producing walls but not connected to the plasma membrane, and (C) cells producing minute amounts of wall material, according Davies and Plaskitt (1981).

**Figure 5.**
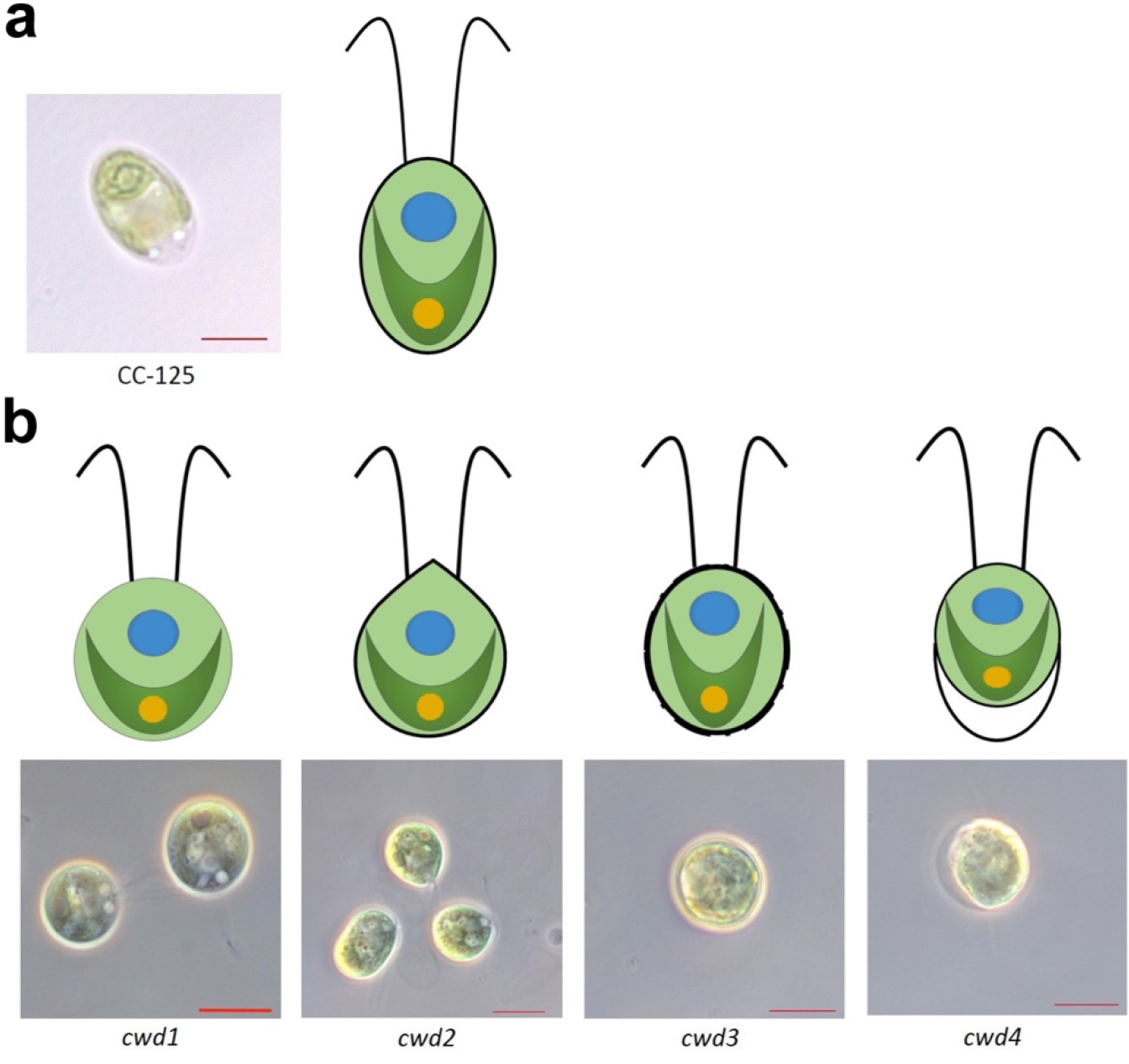
Phenotypic characteristics of wild-type and selected cell wall-defective mutant lines. (**a**) Left image, micrograph of wildtype cell (labeled below), scale bar = 5μm; right image, illustration of wildtype cell showing typical morphology. (**b**) Upper row, artistic renderings of *cwd* mutant cells illustrating variations in cell wall or lack of cell wall; bottom row, corresponding light micrographs of representative cell wall defective mutants (labeled below), scale bars = 10μm.

RT-qPCR analysis showed that all *cwd* mutants expressed a much higher level of CW genes compared to the wild-type before g-lysin treatment (Figure 6). When compared to the upregulated expression levels of the g-lysin-treated wild-type strain, untreated *cwd2, cwd3,* and *cwd4* cells exhibited comparable or higher expression of *PHC19* and *GAS28*, whereas *cwd1* was found to show four to five times less expression of *PHC19* and *GAS28* than the other *cwd* mutants (Figure 6b). *cwd2* and *cwd4*, which exhibited the highest CW gene expression, showed no further up-regulation following g-lysin treatment, suggesting that the g-lysin-induced signaling was fully activated before g-lysin treatment (Figure 6c, e). On the other hand, *cwd1* with the modest CW gene expression showed a full-scale response comparable to the wild-type following the g-lysin treatment, and *cwd3* with the comparable CW gene activation to the g-lysin-treated wild-type showed 3.3- and 3.6-fold further up-regulation of *PHC19* and *GAS28* (Figure 6d). This residual response to g-lysin suggests that g-lysin-induced signaling was partially activated in *cwd1* and *cwd3*.

**Figure 6.**
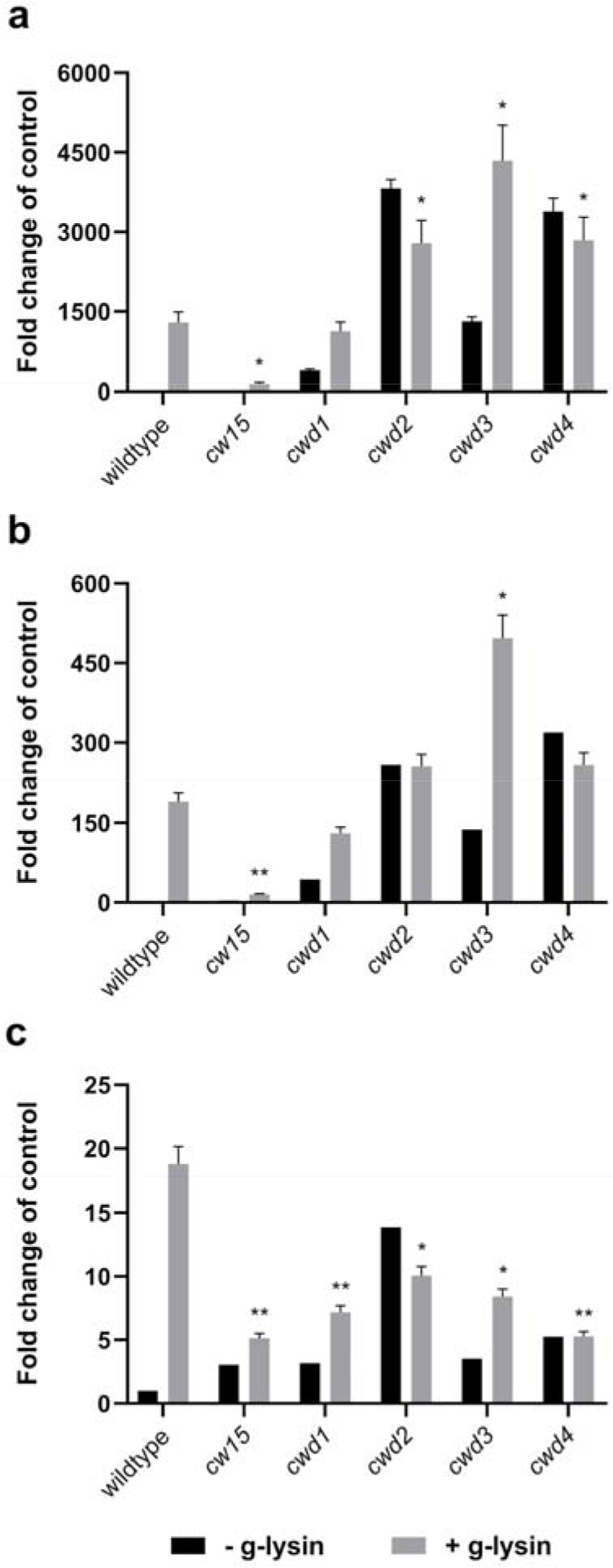
The CW genes are constitutively activated in cell wall defective mutants. (**a-c**) Bar graphs represent the change in gene expression by g-lysin treatment for *PHC19* (a), *GAS28* (b), and *SEC61G* (c) in wildtype (CC-125), *cw15*, *cwd1*, *cwd2*, *cwd3*, and *cwd4*. Untreated gamete control samples represented by black bars, -g-lysin; medium grey bars for g-lysin-treated cells, +g-lysin. Gene expression is quantified in terms of fold change of the untreated wildtype cells. Error bars represent one standard deviation from the mean of biological duplicate samples. Welch’s t-test indicates statistical significance in difference from the g-lysin-treated wildtype at p ≤ 0.05 (*), p ≤ 0.01 (**); α = 0.05.

This gene expression analysis of *cwd* mutants suggests cell wall integrity as a critical factor for the CW gene regulation in *C. reinhardtii*. The result of *cw15*, however, showed an interesting exception. *PHC19* and *GAS28* were found to be expressed higher in untreated *cw15* than in the wild-type, yet at least 30-fold lower than in *cwd1* – showing the lowest CW gene expression among the *cwd* mutants (Figure 6a, b). g-lysin treatment upregulated *PHC19* and *GAS28,* but at a modest level, less than ten-fold in *cw15*.

We reasoned that the low CW gene expression of *cw15* even with g-lysin treatment might be due to its long-term adaptation to its cell wall-less condition where constant CW gene expression becomes wasteful. It is, therefore, conceivable that *cw15* may have accumulated a mutation preventing CW gene expression. To learn about whether such a suppressor mutation exists in *cw15*, sexual-recombinant progeny were generated by mating *cw15* with a cell wall-intact strain transformed with the *PHC19*-luciferase construct (PHC19-2 in Figure 2). A consistent 2:2 segregation of the cell wall defect among the progeny indicates that a single *cw15* mutation is likely responsible for the cell wall defect in the recombinant progeny (data not shown). We selected one tetrad in which two progeny with the *PHC19* reporter are distinguished by their cell wall phenotypes: one with intact cell walls as a *CW15* strain and the other with defective cell walls as a *cw15* strain. Luciferase activity of the *CW15* progeny showed a 4.9-fold increase by the g-lysin treatment, whereas the *cw15* progeny showed constitutive reporter activity at the similarly high level of the g-lysin-treated *CW15* strain (Supplemental Table S2). Upregulated *PHC19* and *GAS28* gene expression ascertained the recovery of CW gene expression in the selected *cw15* progeny (Supplementary Figure S3). These results indicate that compromises in cell wall integrity trigger the upregulation of CW genes in *C. reinhardtii*.

## Discussion

This study focused on how cells sense ‘nakedness’ to rebuild their wall. By analyzing CW genes activated by cell wall removal and imperfect cell walls, we propose a signaling pathway by which physical integrity of the cell wall turns on the CW genes required for cell wall assembly/remodeling in *C. reinhardtii*.

The cell wall is the outermost layer that plays a significant role in water balance by protecting cells from bursting in hypotonic conditions like the freshwater environment (and natural habitat of *C. reinhardtii*). It is, therefore, expected that the removal of the cell wall would have an impact on osmoregulation. When wild-type CC-125 cells were transferred to media of varying osmolarity, we observed little change in CW gene expression level (Supplementary Figure S2). In contrast, we observed 72%-92% reduction of the g-lysin-induced CW gene activation in iso-or hypertonic conditions (Figure 4). In the iso- and hypertonic media, contractile vacuole cycling was not observed, indicating that water influx was not significant. In the absence of water influx, cells may survive even if the cell wall is not perfect or absent. It is, therefore, logical that building an intact cell wall may become less critical under iso-or hypertonic condition.

The second hypothesis tested is the involvement of cell wall integrity signaling that senses structural defects in the cell wall. In comparison to the g-lysin-treated naked cells, full-scale CW gene activation was observed in *cwd2* (with an enclosed cell wall) and in *cwd4* where cells are either enclosed or detached with the cell wall but still maintain contacts with the wall (Figure 6). Whereas, *cwd1* (with little cell wall) showed 20%-30% of the full activated CW gene expression level with further upregulation by g-lysin treatment (Figure 6). These results raise critical aspects of the signaling induced by g-lysin treatment. First, cells lacking a standard cell wall continuously attempt to rebuild their cell wall as indicated by constitutive activation of CW genes. Secondly, naked cells do not induce full-scale CW gene activation, and prepared g-lysin lysates containing digested cell wall material further up-regulate the CW genes. Collectively our findings suggest that the activation of CW genes is likely triggered by cell surface receptors serving two roles: 1) to monitor tension built up between the cell wall and turgid cells that is lost in naked cells and iso- or hyperosmotic conditions, and 2) to sense a cell wall component or specific structure of the typical cell wall that are exposed to the cell surface in cell wall defective mutants and by g-lysin treatment.

Cell wall integrity signaling has been extensively studied in yeast and more recently in plants. In *S. cerevisiae*, the Wsc1 and Mid2 serve as the primary sensor of the yeast cell wall integrity, both of which are a single-transmembrane domain protein with a stretchable *O*-mannosylated Ser/Thr-rich extracellular domain upon mechanical stresses on the cell wall^23,24^. These sensors are coupled with a small G-protein Rho1 for downstream signaling involving Pkc1 and the MAP kinase cascade^25^. Interestingly, this Rho1-Pkc1-mediated signaling and osmotic stress share Skn7, a two-component response regulator, as a downstream transcriptional regulator^26^. Based on the strong down-regulation of CW gene activation under iso- and hypotonic conditions, similar crosstalk between the cell wall integrity signaling and osmotic stress signaling may also exist in *C. reinhardtii*.

In plants, the *Catharanthus roseus* receptor-like kinase like (*Cr*RLKL) family such as ANX and FER have an extracellular domain containing one or two malectin-homology domains that can bind to di-glucose, whose signaling is known to regulate various plant wall features such as stiffening^27,28,29^. BRI-dependent brassinosteroid signaling is another critical player in cell wall integrity signaling, which leads to the activation of cell wall loosening genes^30^. Although no homologs to *Cr*RLKLs or BR-signaling components are found in *C. reinhardtii*, surface receptors localized in the plasma membrane have been reported, including the sex-agglutinins that mediate gamete recognition during mating^31,32^, and the elusive receptor for the sex-inducing pheromone studied in *Volvox carteri*, a close relative of *C. reinhardtii*^33,34^. Based on the distinct structure and composition of *C. reinhardtii* cell walls, we would not expect to find homologs to the known sensors in yeast and plants.

A plausible approach to identify the molecular components of the unknown cell wall integrity signaling in *C. reinhardtii* is to look for mutations that eliminate the constitutive activation of CW genes in cell wall defective mutants, using a forward genetic approach. Such a suppressor screen can be performed, using a reporter system probing the CW gene activation such as fluorescent proteins or colorimetric assays. Molecular details of cell wall integrity signaling components in *C. reinhardtii* will invite comparative studies that ask whether cell wall signaling is conserved between plants and *C.reinhardtii* despite their distinct cell wall composition and organization.

## Methods

### Strains and culture conditions

*C. reinhardtii* strains, CC-125 (wild-type, *mt+*), CC-621 (*mt*-), CC-2663 (*nic7*; *mt*-) and CC-3491 (*cw15*, *mt*-) were obtained from the Chlamydomonas Resource Center (www.chlamydollection.org). All *C. reinhardtii* strains were maintained and cultured under medium light (50 μmol photons m^−2^ s^−1^) at 23°C in Tris-acetate-phosphate (TAP) medium1 (Harris, 1989) solidified with 1.5% Bacto agar. *cwd1, cwd2, cwd3, and cwd4* were isolated from mutagenized population of JL28 (*nic7*; *mt*-) that was generated by mating between CC-125 and CC-2663 (*nic7*; *mt*-). Nitrogen-free TAP medium was prepared by omitting nitrogen from TAP medium. Liquid media with varying osmolarities were made by either adding amounts of sucrose to liquid TAP media or by diluting the media with water. ½ TAP, 1:1 water to media; TAP+S, 60 mM sucrose; TAP+SS, 120 mM sucrose.

### Gametogenesis

Seven-day-old cells grown on TAP plates were harvested and suspended in nitrogen-free TAP (NF-TAP), then counted and normalized to a concentration of 5 × 10^7^ cells/mL. Suspended cells were incubated under high light (200 μmol photons m^−2^ s^−1^) for a minimum of 3 hours to induce gametogenesis. Sufficient gametogenesis was determined by a mating efficiency analysis as per the method described in Hoffman and Goodenough^35^, with 80% mating efficiency being the acceptable lower limit before continuing the experiment.

### Isolation of cell wall defective mutants by insertional mutagenesis

To isolate cell wall defective mutants, we followed the mutant screen using the method described by Davies and Plaskitt^20^. Abnormal colony morphology was screened by scanning plate cultures on TAP medium solidified with 2% Bacto agar under a dissecting microscope (S8APO, Leica) in a mutant pool generated by insertional mutagenesis. Putative mutant colonies were resuspended in liquid TAP medium and tested for the NP-40 sensitivity (detailed in the methods section). Mutants displaying >50% NP-40 sensitivity were selected as cell wall defective (*cwd*). Insertional mutagenesis was performed by the glass bead-assisted transformation using nicotinamide-requiring mutant cells (*nic7*, mt-) as described^36^. The plasmid used in the insertional mutagenesis was prepared by adding pHsp70A/RbcS2-AphVIII^37^ in pNic7.9^38^. The plasmid was linearized by *Eco*RI that cleaves between the pHsp70/RbsC2 promoter and the open reading frame of the AphVIII.

### g-lysin and CHX treatment

g-lysin extract was prepared as described^13^. Prepared g-lysin extract was then frozen at −80°C until use. For g-lysin treatment, suspended cells were mixed with an equal volume of thawed g-lysin extract for 1 hour to ensure cell wall removal. g-lysin efficiency was determined by NP40 sensitivity test (see “NP40 sensitivity testing” in Materials and methods). Untreated control samples were mixed with an equal volume medium to maintain equal cell concentrations across samples. Cells were then incubated for 1 hour before harvesting RNA.

For CHX pretreatment, suspended cells were mixed with a small volume of 10 mg per mL CHX stock to a final concentration of 10 µg/mL, then incubated for 45 minutes. Following the pretreatment, some cells were treated further with an equal volume of thawed g-lysin extract. Cells were then incubated for one hour before harvesting RNA.

### Reverse transcription and quantitative PCR (RT-qPCR)

Total RNA extraction, cDNA synthesis, and qPCR reactions were carried out essentially as described^19^. In brief, each qPCR run had technical duplicate samples to generate average quantification cycle (Cq) data per run. Relative expression levels in each cDNA sample were normalized to the *RACK1* reference gene under the same conditions. Relative expression was calculated by the method described in Pfaffl^39^, which accounts for difference in PCR primer efficiency for the different transcript targets. Two or three biological replicates were averaged for quantitative analysis. Welch’s t-test was applied to the analyzed expression data to check for statistical significance between treatment conditions. Primer sequences are presented in Supplementary Table S3.

### Luciferase activity assays

To induce promoter-reporter activity, *promoter-luciferase*-transformed lines were subjected to treatment conditions. Secreted *Gaussian* luciferase enzyme (from *Gaussia princeps, codon-optimized for C. reinhardtii*^40^) was collected from samples with equal cell concentrations by aliquoting 100 μL of sample into 1.5mL centrifuge tubes and centrifuging for 3 minutes at 6,000 g. 40 μL of supernatant was then transferred from each sample to PCR strip tubes and mixed with 10 μL *Renilla* luciferase lysis buffer (Promega). Prepared samples were then stored at −20°C until luciferase assays were run. Samples for luciferase assay were prepared by mixing 25 μL luciferase assay reagent (luciferase assay buffer + 1x final concentration luciferase assay substrate) (Promega) with 5 μL of thawed samples in a 384-well microtiter plate. The relative amount of luciferase enzyme in each sample was then quantified by the amount of luminescence detected by a BioTek Synergy 2 microplate reader.

### NP-40 sensitivity testing

20 μL of suspended cells were mixed with 20 μL 0.2% NP-40 “Tergitol” detergent (Sigma) in a 1.5mL tube. Another 20 μL of suspended cells were simply diluted with media as an untreated control. 10 μL of mixture and 10 μL of untreated cells were loaded onto either side of a hemocytometer and visualized using brightfield microscopy with 100x total magnification. Both treated and untreated cells were counted to compare cell concentrations and determine the number of cells that had burst from the NP-40 treatment. This detergent disrupts the lipid-based membrane and will therefore cause any cells without an intact cell wall to lyse. A simple ‘NP-40 sensitivity’ value was then calculated by dividing the untreated cell concentration by the treated cell concentration and multiplying by 100%. For example, if all cells in a sample treated with NP-40 burst, the NP-40 sensitivity would be 100%.

### Contractile vacuole visualization and timing

To ascertain osmotic conditions of the media used in experiments, we measured the CV cycle (contractile vacuole cycling times) in the wild-type and a cell wall-less strain, *cw15*, under various osmotic conditions as defined in Komsic-Buchmann et al.^41^, either hypotonic media, half-diluted standard medium TAP (½ TAP, 32 mOsm/L) and the standard TAP (64 mOsm/L), or hypertonic media, the TAP supplemented with sorbitol (TAP+S, 144 mOsm/L; TAP +SS, 204 mOsm/L) and incubated for an hour. 8 μL of cell suspension was loaded onto glass slides and cells were viewed under 1000X total magnification using Zeiss Axioscope A1. The CV cycle was measured manually by timing consecutive systole and diastole cycles of a single contractile vacuole per cell. A minimum of 3 cells per cell line per osmotic condition were measured.

We confirmed that increasing osmolarity in the media extended the CV cycles, indicating slow water influx. This trend was found in all the cell lines tested (Supplemental Table S1). The CV cycles were observed in the TAP+S condition but none in the TAP+SS condition, indicating that cells have their osmolarity between 144 and 204 mOsm per L, which agrees with the published cytosolic osmolarity of ~171 mOsm per L^42^. The *cw15* cells showed much longer contractile vacuole cycle times, nearly double the length of the wild-type cycles. In pure water (0 mOsm/L), the wild-type cells showed a contractile vacuole cycle time at the average of 9.56s, whereas the cw15 cells exhibited an average time of 17.66s. The correlation between the cell surface area and the CV cycle was previously noted^41^.

## Acknowledgments

This work was supported by Discovery Grant 418471-12 from the Natural Sciences and Engineering Research Council (NSERC) (to J.-H.L.), by the Korea Carbon Concentration and Sequestration Research and Development Center (KCRC), Korean Ministry of Science, grant no. 2016M1A8A1925345 (to J.-H.L.). We thank Dr. Sunjoo Joo for critical reading of the manuscript and Jenny Lee for technical assistance.

## Author contribution

EC, and J-HL designed the research; EC, DT, NCS, TK, MHW, and J-HL performed the experiments. EC and J-HL analyzed the data. EC and J-HL wrote the article. All authors reviewed the manuscript.

## Additional information

The authors declare no conflict of interest.

## Supplementary information

**Supplementary Text S1.** Osmoregulation in *Chlamydomonas reinhardtii.*

**Supplementary Table S1.** The CV period (contractile vacuole cycling times) in five different media of varying osmotic conditions.

**Supplementary Table S2.** *PHC19* promoter-driven luciferase activity of cw15 progeny in response to g-lysin treatment.

**Supplementary Table S3.** Primers used in this study.

**Supplementary Figure S1.** Hypothetical signaling mechanisms for the g-lysin-induced responses.

**Supplementary Figure S2.** Transcript expression suggests modest cellular response to changing osmotic environment.

**Supplementary Figure S3.** *cw15* progeny showed constitutive CW gene expression.

**Supplementary Figure S4.** Diagrammatic representation of osmoregulation in *C. reinhardtii* cells.

